# Dentate spikes comprise a continuum of relative input strength from the lateral and medial entorhinal cortex

**DOI:** 10.1101/2025.10.27.684857

**Authors:** Gergely Tarcsay, Rajat Saxena, Royston Long, Justin L. Shobe, Bruce L. McNaughton, Laura A. Ewell

**Affiliations:** Department of Anatomy & Neurobiology, University of California, Irvine, CA 92697; Department of Neurobiology and Behavior, University of California, Irvine, CA 92697; Canadian Centre for Behavioural Neuroscience, The University of Lethbridge, Lethbridge, Alberta T1K 3M4, Canada; Kavli Institute for Systems Neuroscience and Centre for Algorithms in the Cortex, Norwegian University of Science and Technology, Trondheim, Norway

## Abstract

Hippocampal dentate spikes (DS) are synchronous population events generated in the dentate gyrus thought to support memory processing. DS are classified into type 1 (DS1) and type 2 (DS2) based on current sinks in the outer or middle molecular layer, where the axons of the lateral or medial entorhinal cortex terminate, respectively. This widely used classification method constrains DS into a bimodal distribution without properly testing whether such constraint is appropriate. Here we utilized silicon probe recordings with high spatial resolution spanning all layers of the dentate gyrus and discovered that the contribution of LEC/MEC inputs to DS follows a continuous distribution. We introduce a third type (DS3) that captures a previously unreported component of the distribution: simultaneous current sinks in the outer/middle molecular layer, suggestive of synchronized activation from the LEC/MEC which could facilitate binding. We verify DS3 in several data sets from independent laboratories and characterize their brain state dependence and neural recruitment.

## INTRODUCTION

The dentate gyrus (DG) network is positioned at the entryway of the hippocampal circuit, transforming cortical signals before transmitting to downstream CA3/CA1 subregions and plays a critical role in memory processing^1–6^. The lateral and medial entorhinal cortices (LEC/MEC) send converging projections to the DG, conveying primarily non-spatial and spatial information respectively^7–12^. Thus, one major function proposed to be performed by the DG is to integrate spatial and non-spatial signals into a single representation, termed as conjunctive coding^13–17^. Behavioral studies have shown that DG is essential in memory tasks requiring the associations between objects, context and space^18–21^. Multimodal representations in DG principal cells further support the existence of a conjunctive code, however, whether multimodality is constructed in the DG proper or inherited from cortex remains poorly understood^22,23^.

There are two somewhat contradictory major electrophysiological signatures of the DG. First, the DG is known for its relative low firing ‘quietness’ compared to upstream and downstream regions^24^. There is a large expansion in cell number from the input layers (MEC and LEC) to the DG granule cell layer^25^. The large DG neural population combined with powerful circuit inhibition mediates a winner-take-all coding scheme which results in an extremely sparse activation of DG granule cells^26–29^. Such sparsity is thought to endow the DG with the ability to efficiently pattern separate incoming cortical inputs^30^. The quiet of the DG is punctuated with the second signature: dentate spikes (DS). DS are large amplitude positive deflections in the hilar local field potential (LFP) accompanied by synchronous population bursts^31–33^. DS are prominent during immobility and slow-wave sleep and are proposed to support associative learning and to facilitate flexible memory storage^31,33–38^, however much remains to be learned about these highly synchronous events in an otherwise quiet neural structure.

Most studies classify DS into two distinct populations based on the pattern of activation from the entorhinal cortex. DS type 1 (DS1) comprise a current sink in the outer molecular layer (LEC projection) and DS type 2 (DS2) comprise a current sink in the middle molecular layer (MEC projection)^31,33,38–41^. Classification of DS into these two distinct types suggests that during DS, spatial and non-spatial inputs are processed separately. In contrast, recent findings of EC to DG signaling during ongoing gamma oscillations suggest that both input streams are co-active when behavior tasks engage multiple modalities, such as during object-place memory^42^. Here we investigate whether the distribution of DS is truly bimodal. Recording with high-density silicon probes in freely moving mice, we find that DS comprise a continuum of inputs from the MEC and LEC. We define a new class, DS type 3 (DS3), that are accompanied by synchronous current sinks in the outer and middle molecular layers. We confirm that DS3 are present in data sets of multiple laboratories, including those acquired from head-fixed mice. DS3 show brain state dependence with moderate rates during quiet wakefulness and active exploration, and near complete suppression during slow-wave sleep. Furthermore, the majority of recorded DG neurons were recruited during DS3, highlighting their relevance in impacting downstream hippocampal processing.

## RESULTS

### Classification of dentate spikes by input patterns from the entorhinal cortex

We performed high-density silicon probe recordings from the DG in awake, freely moving mice (N=12) and detected dentate spikes (DS) in the hilar LFP (Figure 1A, Figure S1A). To localize the outer molecular layer (oml) and middle molecular layer (mml) where the LEC and MEC terminate, respectively, we performed current-source density (CSD) analysis on each putative DS (Figure 1B; see Methods for details). Next, we identified the channels where the deepest current sinks of either DS1 or DS2 were observed and considered those channels as the locations of the oml and mml respectively, similar to^43^ (Figure 1B; Figure S1B). To investigate the relative contributions of LEC/MEC inputs to DS, we visualized the magnitude of current sinks for both the oml and mml for each DS similar to^35^. Strikingly, we did not observe two clusters as would be expected if the distribution included only DS1 and DS2 (Figure 1B, cyan and magenta). Instead, relative oml-mml contributions formed a continuum, with a large subset of events having contributions from the two inputs (Figure 1B, grey). To quantify the contribution of LEC and MEC to the DS with two current sinks (N = 2982 of 5096), we calculated a score that reflected the relative current sink magnitudes [(oml-mml) / (oml+mml)]. We found a continuous distribution of relative current sinks that on average was skewed to stronger MEC contribution (Figure 1C, Figure S1C). We thus define a novel type of dentate spike, DS type 3 (DS3) that is accompanied by simultaneous oml and mml current sinks (Figure 1C, D, scores ranging from −0.5 to 0.5; Figure S1D, E). We further found that the magnitudes of the current sinks were generally smaller for DS3, potentially highlighting the importance of the synchronous dual input for recruiting populations of cells which will be addressed in later figures. (Figure S1D, mml sink DS2: 0.1 ± 0.02 mean ± s.d vs. DS3: 0.03 ± 0.01, paired t-test with post hoc Bonferroni correction; p < 0.0001, t(19) = −18.4, oml sink DS1: 0.05 ± 0.01 vs. DS3: 0.03 ± 0.01, p < 0.0001, t(19) = −7.8,, N = 12 freely moving mice, N = 8 head-fixed mice).

**Figure 1.**
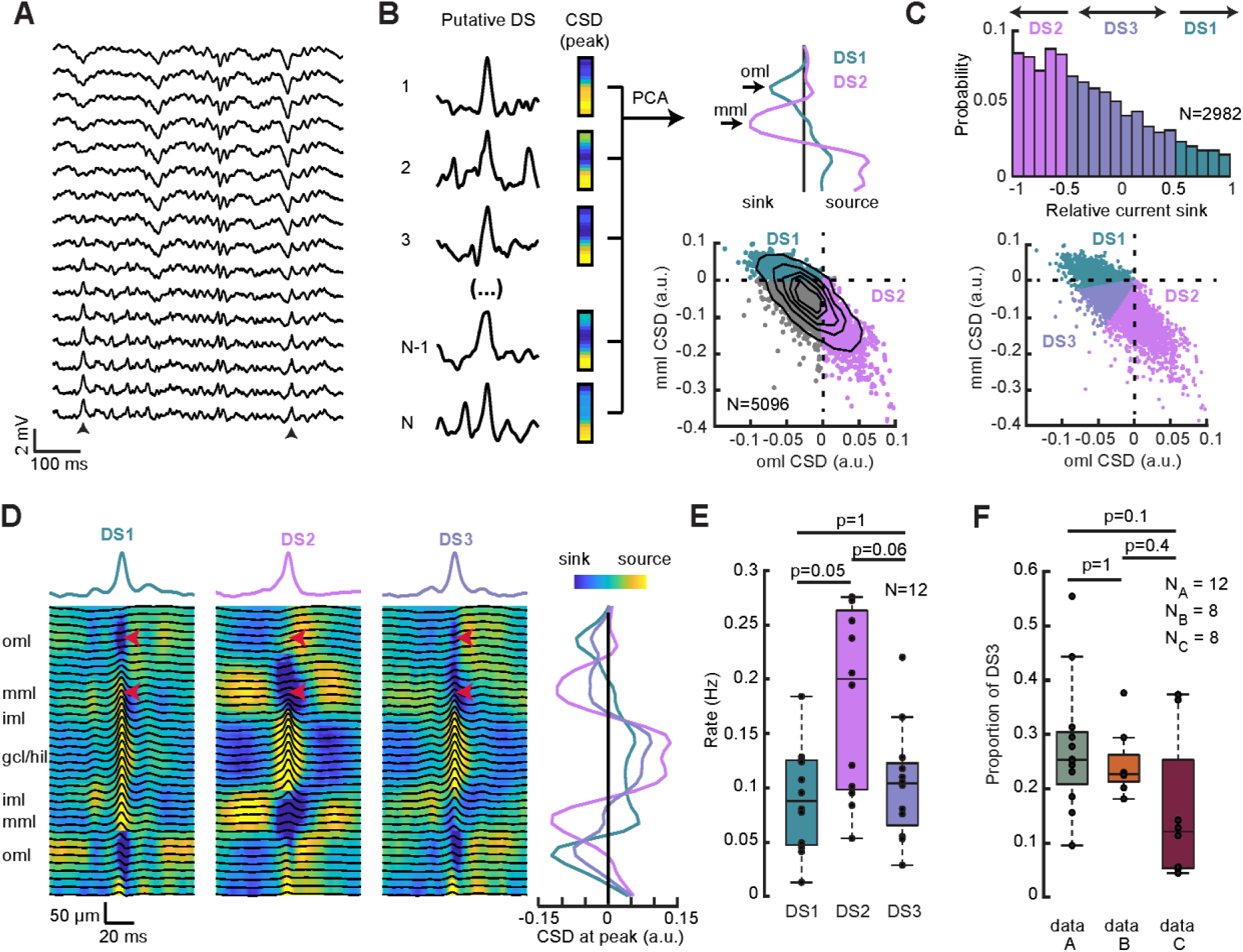
Classification of dentate spikes into 3 types based on CSD profile. (**A**) Silicon probe recordings spanning from the hilus to the fissure with 25 µm spacing between channels. Example shows two putative DS largest on hilar channels (black arrows). (**B**) Schematics of the DS classification algorithm. Current-source density was calculated across channels for each putative DS and PCA was performed on them (left). Location of deepest sinks associated with DS1 and DS2 were used to represent the oml and mml and DS were mapped into the oml-mml state space (right, top). DS1 are marked in cyan, DS2 are marked in magenta. DS associated with sinks in both layers are marked in gray. (**C**) Distribution of relative current sink of DS with sink both in oml and mml. DS3 (purple) were defined if the relative current sink was between −0.5 and 0.5. (**D**) Average waveform and CSD for each DS type from one mouse. Red arrows mark the outer and middle molecular layer. (**E**) Rates of the 3 DS types during (N=12). DS2 were significantly higher than DS1 and a trend was observed compared to DS3. (**F**) Proportion of DS3 in 3 distinct data sets.

We found that DS3 and DS1 occurred at similar rates, while DS2 rates were elevated in comparison, though rates were only significantly different for DS1 vs. DS2. (Figure 1E, repeated-measure ANOVA with post hoc Bonferroni tests, p = 0.007; F(2,22) = 6.2; p_DS1-DS2_ = 0.05; p_DS1-DS3_ = 1; p_DS2-DS3_ = 0.06; DS1: 0.09 ± 0.05 Hz, DS2: 0.18 ± 0.08 Hz, DS3: 0.10 ± 0.05 Hz, mean ± s.d.). We next asked whether DS3 are unique to our experimental paradigm (data A). We extended our analysis to include DS in awake, head-fixed mice (data B, N=8) and in a publicly available dataset^40^ (data C, N = 8). Length of the recording sessions in data C were unknown, therefore we calculated the proportion that were DS3 and found it to be similar across all data sets (Figure 1F, one-way ANOVA with post hoc Bonferroni tests; p = 0.1, F(2,25) = 2.6; p_A-B_ = 1, p_A-C_ = 0.1, p_B-C_ = 0.4, data A: 0.28 ± 0.12, data B: 0.25 ± 0.06, data C: 0.16 ± 0.13; mean ± s.d.). As data C provided pre-detected putative DS events utilizing a more conservative peak detection algorithm described in^40^, this result shows that the observation of DS3 is robust and independent of the detection algorithm.

### Behavioral brain-state dependency of dentate spikes

Previous works showed that DS rates are brain-state dependent and most prominent during slow-wave sleep (SWS) and quiet wakefulness (QW), while reports about DS rates during locomotion are controversial^34,38,39,41^. First, we investigated DS in freely moving mice (N = 12) and found that a substantial portion occurred while mice were running (Figure 2A; DS1: 0.05 ± 0.03 Hz, DS2: 0.10 ± 0.08 Hz, DS3: 0.06 ± 0.05 Hz, mean ± s.d.). Run-related DS rates did not differ from each other for the three DS types (Figure 2B, repeated-measure ANOVA with post hoc Bonferroni tests, p = 0.14; F(2,22) = 2.14; p_DS1-DS2_ = 0.4; p_DS1-DS3_ = 1; p_DS2-DS3_=0.4). We further found that run-related DS rates were significantly lower compared to during QW for DS1 and DS2 and a similar trend was observed for DS3 (Figure 2C; paired t-test with Bonferroni correction, DS1: t(11) = 4, p = 0.006; DS2: t(11) = 4, p = 0.006; DS3: t(11) = 2.6, p = 0.07). DS2 have been linked to transitions from offline to online behavioral state^38,44^. We therefore tested whether DS were associated with changes in running speed. We found that the occurrence of DS1 was associated with slowdowns, however DS2 and DS3 occurred at constant speeds (Figure S2A). To further investigate this observation, we asked when during an individual running epoch DS occur (see Methods for details) and found that the DS1 distribution significantly deviated from a uniform distribution and is more likely to occur at the end of the running epoch (i.e. at transition between running and immobility), while DS2 and DS3 occurred uniformly (Figure S2B, Chi-square goodness-of-fit with Bonferroni correction, DS1: p < 0.0001, χ^2^ = 47.5, df = 9; DS2: p = 1, χ^2^ = 8.5, df = 9; DS3: p = 0.06, χ^2^=19.8, df = 9).

**Figure 2.**
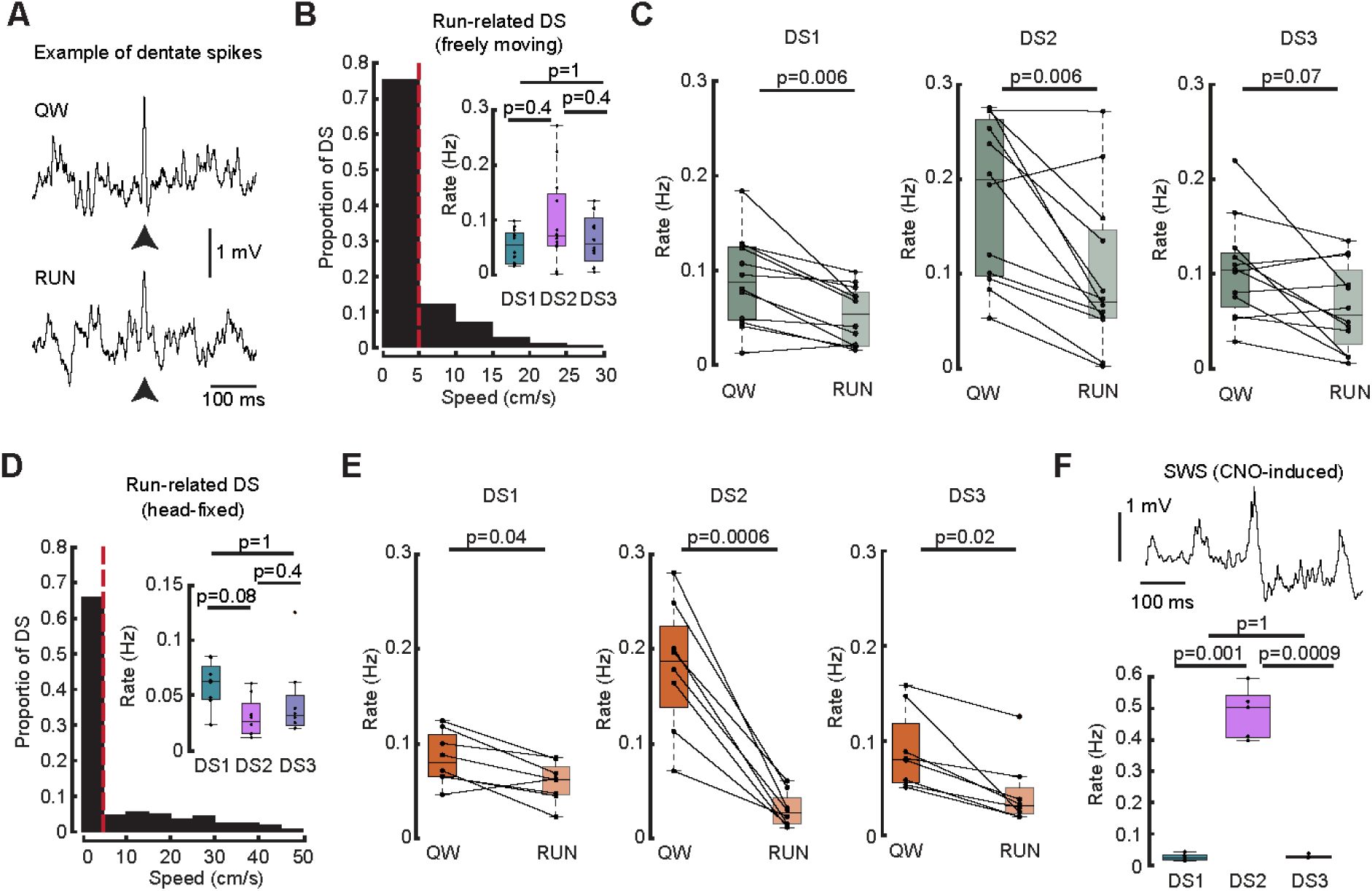
Behavioral brain-state dependency of dentate spikes. (**A**) Example dentate spikes (black arrow) during quiet wakefulness (QW) and during locomotion (RUN). (**B**) Proportion of DS as a function of speed in freely moving animals. RUN-related (> 5 cm/s) DS rates of the three types did not differ from each other. (**C**) Comparison of DS rates between QW and RUN states for DS1 (left), DS2 (middle) and DS3 (right) in freely moving mice. (**D**) Same as in (B), but for head-fixed mice. (**E**) Same as in (C), but for head-fixed mice.

We next asked whether run-related DS are present in behaving head-fixed mice (N=8) in which the extent of sensory inputs may be decreased during locomotion compared to freely moving behavior. Again, run-related DS rates were similar across types (Figure 2D, repeated-measure ANOVA with post hoc Bonferroni tests, p = 0.08; F(2,14) = 3.1; p_DS1-DS2_ = 0.4; p_DS1-DS3_ = 1; p_DS2-DS3_ = 0.4; DS1: 0.06 ± 0.02 Hz, DS2: 0.03 ± 0.02 Hz, DS3: 0.04 ± 0.04 Hz). Similarly to the freely moving case, DS rates during running were lower compared to during head-fixed QW DS rates (Figure 2E, paired t-test with Bonferroni correction, DS1: t(7) = 3.3, p = 0.04; DS2: t(7) = 7, p=0.0006; DS3: t(7) = 3.9, p = 0.02). Furthermore, in head-fixed animals we observed constant running speeds in the second surrounding each DS for all types (Figure S2C). DS1 and DS2 occurred uniformly during individual running epochs, while DS3 occurred at earlier times (Figure S2D, Chi-square goodness-of-fit with Bonferroni correction, DS1: p = 1, χ^2^ = 9.4, df = 9; DS2: p = 0.22, χ^2^ = 15.7, df = 9; DS3: p = 0.03, χ^2^ = 21.9, df = 9). DS rates between freely moving and head-fixed animals did not differ from each other during QW (Figure S2E, unpaired t-test with Bonferroni correction, DS1: p = 1, t(18) = 0.2; DS2: p = 1, t(18) = −0.04, DS3: p = 1, t(18) = 0.6) or during running (Figure S2F, unpaired t-test with post hoc Bonferroni correction, DS1: t(18) = −0.6 p = 1; DS2: t(18) = 2.3, p = 0.1; DS3: t(18) = 1.1, p = 0.9). In summary, DS are more frequent during immobility, however, moderate rates are observed during locomotion, thus potentially contributing to online hippocampal memory processing.

Finally, we asked how rates of distinct DS types are impacted by CNO-induced slow wave sleep (SWS, N = 5, see Methods for details)^45^. We found that DS1 and DS3 rates were significantly decreased compared to DS2 rates (Figure 2F, repeated-measure ANOVA with post hoc Bonferroni tests; p < 0.0001, F(2,8) = 132.8, p_DS1-DS2_ = 0.001; p_DS1-DS3_ = 1; p_DS2-DS3_ = 0.0009; DS1: 0.03 ± 0.01 Hz; DS2: 0.49 ± 0.08 Hz; DS3: 0.03 ± 0.01 Hz, mean ± sd), suggesting that the EC to DG communication during DS is primarily driven by the MEC in induced SWS. These results suggest that MEC input only is not sufficient to observe DS3, further supporting that DS3 are associated with LEC excitation as well.

### Cell recruitment during dentate spikes

To determine how DS impact signal flow between the entorhinal cortex and the hippocampus, we recorded the activity of excitatory principal cells (PCs, N=176) and narrow waveform interneurons (INs, N=86) of the DG in freely moving and head-fixed mice during QW sessions, and asked whether DG cells are modulated by the different DS types. To quantify how strongly neurons were recruited, we created peri-event time histograms for each cell around DS peaks and averaged the cell activity across events in each 1 ms temporal bin, similar to McHugh et al^33^. The resulting trace was z-scored and the maximum value was considered as the activity score (Figure 3A,B, see Methods). We found that the majority of the neurons (85.9 %) were significantly modulated (activity score > 2.58, corresponding to p < 0.01) by DS, and out of those the largest subset (43.6 %) was active in every DS type, while almost one third of them were active exclusively during DS2 (30.7 %, Figure 3C). Importantly, 57 % of the active cells were modulated by DS3, underscoring their potential relevance in hippocampal processing.

**Figure 3.**
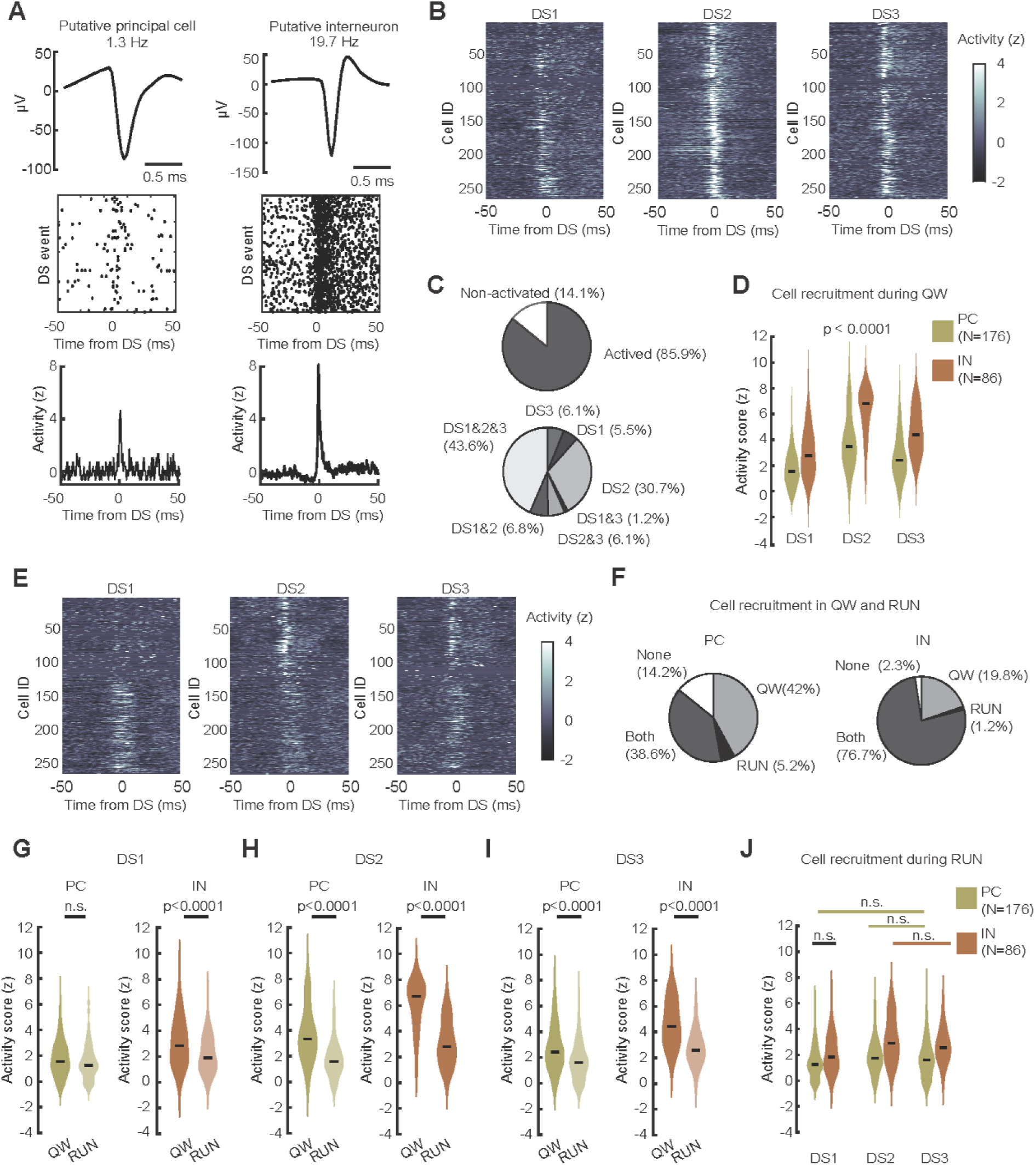
Single-unit recruitment during all types of DS. (**A**) Example waveforms of putative principal cells (PCs) and interneurons (INs) respectively (top). Peri-event time histogram (1 ms wide bins) was created for each cell around DS peaks (middle). Mean firing rate across events were calculated and z-scored across temporal bins (bottom). (**B**) Z-scored activity of the full cell population for the 3 types of DS during QW sessions. (**C**) The majority of the cells were significantly active during DS (activity score > 2.58, top). The largest subpopulation participated in all 3 types of DS, while a third of the cells were modulated by DS2 only (bottom). (**D**) Post hoc comparison of cell modulation in QW state with respect to cell type (PC/IN) and DS type (DS1/2/3). Black line marks the median. Note that each comparison was significant (p < 0.0001, see Table 1 for statistics). (**E**) Cell modulation by the three DS types in RUN state. (**F**) A subset of PCs is modulated during RUN sessions (left) and most of the INs were recruited in RUN (right). (**G**) Post hoc comparison of cell modulation between RUN and QW for PCs (left) and INs (right) during DS1. (**H and I**) Same as in (G), but for DS2 and DS3. (**J**) Post hoc comparison of cell recruitment in RUN state with respect to cell type and DS type. Unlabeled pair-wise comparisons were significant (see Table 1 for more statistics).

**Table 1.**
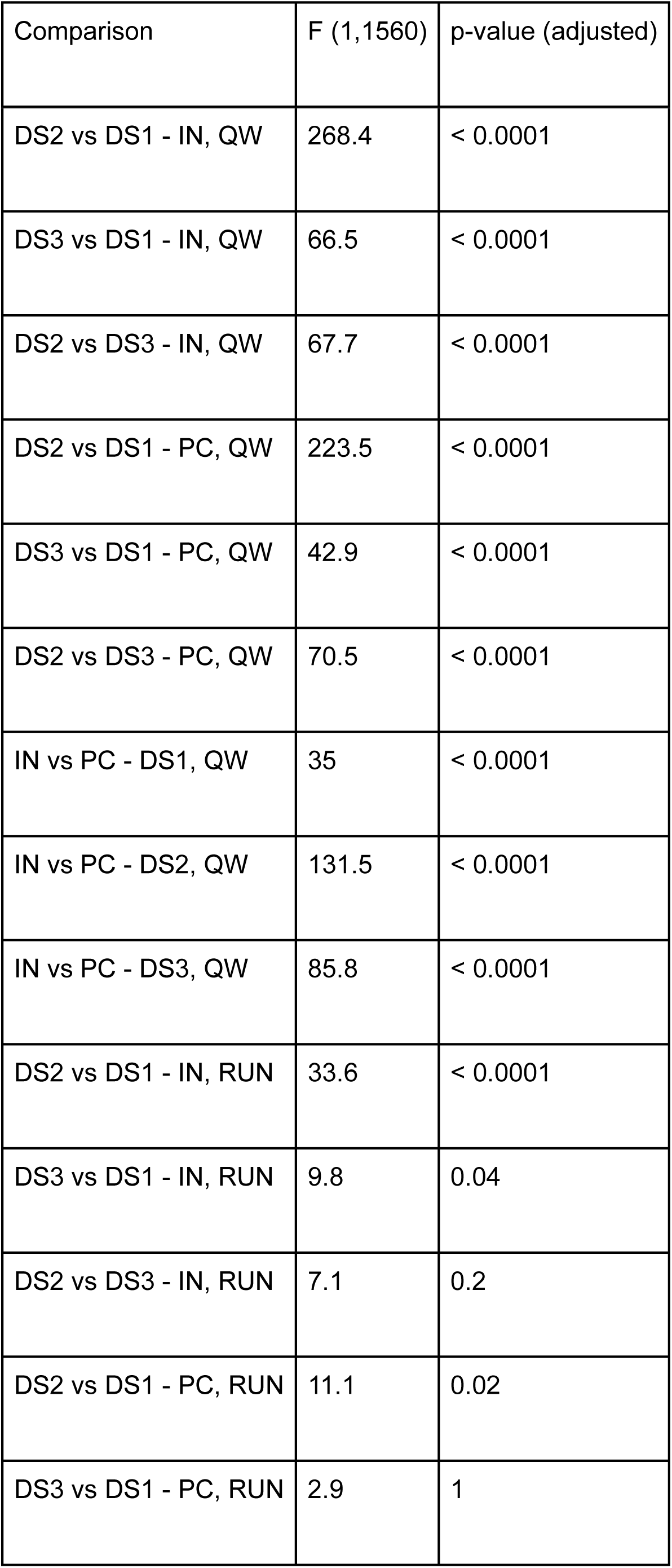
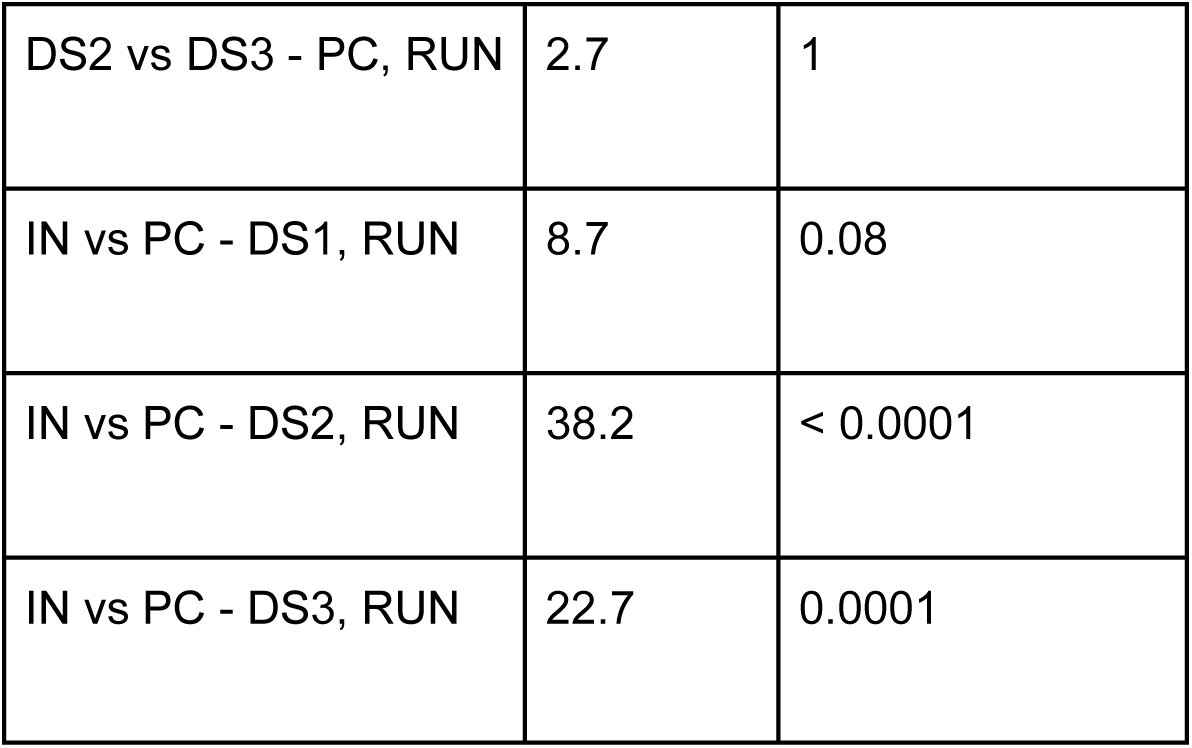
Post hoc multiple comparisons of activity scores for distinct cell types and DS types during QW and during running. Related to. **Figure 3**.

We found that the proportion of active cells during individual DS events was highest for DS2 (Figure S3A; Wilcoxon rank-sum test with post hoc Bonferroni correction ; p_DS1-DS2_ < 0.0001; z = −23; p_DS1-DS3_ < 0.0001; z = −6.7; p_DS2-DS3_ < 0.0001; z = 15.5; DS1: 13 (10; 21) %; DS2: 20 (13;29) %; DS3: 16 (10; 24) %; N = 247 neuron; median (Q1; Q3) percentage of recorded neuron). Individual neurons participated most frequently during DS2 (Figure S3B; Friedman test with post hoc Bonferroni correction; p < 0.0001; χ2(2) = 202.5; p_DS1-DS2_ < 0.0001; p_DS1-DS3_ < 0.0001; p_DS2-DS3_ < 0.0001; DS1: 5.9 (1.7; 17.7) %; N = 2384 DS1; DS2: 11 (3.6; 32) %; N = 4947 DS2; DS3: 8.5 (2.8; 25.4) %; N = 2230 DS3; median (Q1; Q3) percentage of DS).

Next, we constructed a linear mixed-effects model (LME) to ask how cell recruitment is impacted by distinct conditions, such as cell type and DS type (Table S1, see Methods for details). We found a high cell type and DS type specificity of the population activity (ANOVA, cell type: p < 0.0001, F(1,1560) = 35; DS type: p<0.0001, F(2,1560) = 13, cell type x DS type interaction: p < 0.0001, F(2,1560) = 12). Recruitment of INs were significantly higher than PCs for the three types of DS (Figure 3D, post hoc multiple comparison, p < 0.0001 for all pair-wise comparisons, see Table 1 for more statistics; PC: 1.8 ± 1.4, 3.8 ± 21, 2.7 ± 1.7; IN: 3 ± 1.8, 6.2 ± 1.9, 4.6 ± 1.7; for DS1, DS2, DS3 respectively, mean ± s.d. activity score). Furthermore, DS1 exhibited a significantly weaker recruitment of PCs and INs compared to DS2/3s, and DS2 cell modulation was significantly stronger compared to DS3 (Figure 3D). While it may be expected that stronger recruitment is observed when multiple dendritic compartments of granule cells are excited (DS3), compared to when just the dendritic compartment in the middle molecular layer receives inputs (DS2), the magnitude of current sinks accompanied DS3 were significantly lower than the current sinks associated with DS1 and DS2 as discussed previously (Figure S1D). It is noteworthy that despite smaller magnitude sinks, cellular recruitment during DS3 is robust.

### DS cell recruitment during running

DS during locomotion have been reported before^39^, however it is unclear whether these events contribute to neuronal recruitment. Therefore, we calculated the activity score of the same DG neurons during locomotion and found that a moderate portion of PCs (43.8 %) and most INs (77.9 %) were significantly modulated. Similar to during quite wakefulness, we found that the proportion of active cells during individual DS events was highest for DS2 and DS3 (Figure S3A; Wilcoxon rank-sum test with post hoc Bonferroni correction; p_DS1-DS2_ < 0.0001; z = −5.4 ; p_DS1-DS3_ = 0.003; z = −3.3 ; p_DS2-DS3_ = 0.2; z = 1.9; DS1: 20.5 (12.9; 29.4) %; DS2: 23.1 (18; 30.8) %; DS3: 23.1(15.4; 30.8) %; N = 247 neuron; median (Q1; Q3) percentage of neurons) and neurons participated most frequently during DS2 (Figure S3B; Friedman test with post hoc Bonferroni correction; p < 0.0001; χ2(2) = 54.6; p_DS1-DS2_ < 0.0001; p_DS1-DS3_ = 0.0001; p_DS2-DS3_ = 0.007; DS1: 8.3 (2.5; 22.4) %; N = 851 DS1; DS2: 13 (4.7; 35.8) %; N = 345 DS2; DS3: 11.8 (3.6; 32.2) %; N = 396 DS3; median (Q1; Q3) percentage of DS).

We found that brain state significantly modulated cell recruitment in the LME (Table S1; ANOVA, brain state: p < 0.0001, F(1,1560) = 24; brain state x cell type interaction: p = 0.009, F(1,1560) = 7; brain state x DS type interaction: p < 0.0001, F(2,1560) = 28). Cell recruitment was similar for PCs, but not for INs during DS1 between brain states (Figure 3G, PC: p = 0.3, F(1,1560) = 6.1; IN: p<0.0001, F(1,1560) = 24), while for DS2/3 the modulation was significantly higher in QW compared to running both for PCs (Figure 3H-I, DS2: p < 0.0001, F(1,1560) = 198.4; DS3: p < 0.0001, F(1,1560) = 53.8) and INs (DS2: p < 0.0001, F(1,1560) = 240; DS3: p < 0.0001, F(1,1560) = 98.7). We found similar modulation between PCs and INs during DS1, moreover DS3 recruitment did not differ from DS2 recruitment in both cell populations and from DS1 modulation for PCs (Figure 3J, see Table 1 for statistics). Taken together, these results show a moderate cell recruitment during DS that occur during running. Consequently, DS recruit neuronal engagement during online hippocampal processing. Notably, we found that cell modulation during running diverges between freely moving mice (Figure 3E, Neuron IDs 1-119) and head-fixed mice (Neuron IDs 120-262). DS1 recruitment was primarily present in the head-fixed paradigm (Figure S3E; Wilcoxon rank-sum test with post hoc Bonferroni correction, p < 0.0001, z = −7.7; freely moving: 0.96 (0; 1.4); head-fixed: 1.9 (1.2; 3.1); median (Q1; Q3) activity score), while DS2 recruitment was more prominent in the freely moving paradigm (p < 0.0001, z = 5.96; freely moving: 2.8 (1.7; 4.3); head-fixed: 1.65 (1.1; 2.4) and DS3 recruitment was similar in the two paradigms (p = 0.1; z = 2.1; freely moving: 2.1 (0.85; 3.25); head-fixed: 1.7 (1.1; 2.6)). Taken together these findings could suggest that the type of external stimuli experienced regulates cellular recruitment during running related DS.

## DISCUSSION

We found that DS are associated with continuously distributed LEC/MEC relative current input, challenging the current view of two well-separated classes of DS (DS1 and DS2). Accordingly, a subset of DS, referred to as DS3, was accompanied by synchronous current sinks in the oml and mml that were present in multiple datasets. As reported for other DS, DS3 exhibited brain state dependence, such that they occurred during quiet wakefulness and running and were almost completely suppressed during slow wave sleep. Furthermore, we observed strong modulation of DG neuronal populations during DS3 that also showed brain state dependence.

There are several possible mechanisms that could produce a current source density pattern consistent with what we observe during DS3. The first is temporally precise coordination of the LEC and MEC to produce synchronous excitation of the oml/mml. LEC(II) fan cells have been shown to project to MEC(I), indirectly exciting stellate cells in MEC(II), serving as a potential mechanism to elicit DS3^46^. Recent studies further showed excitatory and inhibitory connections from MEC to HPC-projecting LEC(II) cells, extending the options of crosstalk between the two regions^47^. A second possibility is that somatic inhibition of granule cells produces large positive deflection in the hilus and corresponding negative components in the molecular layers^48^. However, we found that DS3 are associated with strong modulation of cells (Figure 3D) with roughly 15 % of recorded cell activated during individual DS3, implying an excitatory drive. We propose it is unlikely that DS3 are driven by inhibitory mechanisms exclusively.

The strong engagement of DG neurons during DS3 suggests that these events may elicit conjunctive coding, binding distinct information arriving from the LEC(II) and MEC(II). It is important to note that recent studies highlight the multimodality of entorhinal cortex, suggesting that binding may occur in multiple steps during memory processing^9,49–51^. The hippocampus is known to be critical in paired-associative learning of objects and locations^52–54^, and perhaps DS3 are the underlying neuronal substrate. Hippocampal-dependent behavioral tasks have been designed to test object-place^55^ and object-place-context associations in rodents^56^, however, the role of DS, and more specifically the role of DS3 in these tasks remains to be tested. Another possibility is that DS3 facilitate LEC signaling and synaptic plasticity with the support of MEC excitatory inputs via dendritic integration in granule cells^57–59^.

DS during locomotion are associated with moderate modulation of DG neuronal population, suggesting that DS can reflect online memory processes during active exploration. One possibility is that these events recruit place cells that correspond to the current location of the animal, similar to previously reported offline to online state transitions^38^. Such activation may reinforce the synaptic connection of temporally co-active neurons, supporting a population code for a given environment^60^. As DS3 may carry spatial and non-spatial information, running-related cell modulation may facilitate the reorganization of hippocampal representation between environments.

## METHODS

### Subjects

All experimental procedures were performed as approved by the Institutional Animal Care and Use Committee at the University of California, Irvine. Experiments in the Ewell lab were using C57BL/6 mice (N=12, Charles River), were single housed and kept in an inverted 12:12 hour light/dark cycle, with a temperature of (22±2) °C and relative humidity of (55±10) %. Food and water were available ad libitum, except during behavioral experiments when mice received a 2 % citric acid (CA) water replacement. Weight was monitored and maintained above 85% of the original weight during CA water replacement. Experiments in the McNaughton lab were using B6129SF1/J mice (N = 8, Jackson Labs) and Vgat-Cre transgenic mice (N = 5, Slc32a1^tm2(cre)Lowl^, Jackson Labs) and were housed on a 12:12 hour light/dark cycle.

### Surgical procedure

3-5 months old mice were anaesthetized with 1-2% isoflurane and placed into a stereotaxic frame (David Kopf Instruments). Buprenorphine (0.1 mg/kg) or Carprofen (5mg/kg) was injected to minimize discomfort and mice were kept on a heating pad to maintain body temperature. For chronic freely moving experiments, mice were implanted with a 64-channel high-density linear probe (Neuronexus, H64LP, single shank, 20 μm vertical spacing or Cambridge Neurotech, H2 ASSY-156, double-shank 25 μm vertical spacing) in the cortex above the right hippocampus (AP = −2.1 mm, ML = 1.45 mm from Bregma, DV = −0.9 mm from dura). The craniotomy was covered with Dura-Gel (Cambridge Neurotech) and the implant was shielded by a mouse cap (3Dneuro). Upon surgery, the probe was manually lowered until the infrapyramidal blade of the DG was reached, across 7-12 days. For acute head-fixed experiments, mice were implanted with stainless steel headbars and mice used for the slow-wave sleep analysis were further injected bilaterally with pAAV2-hSyn-DIO-hM3D(Gq)-mCherry virus (Addgene, 100-125 nl, 30-40 nl/min) into the parafacial zone to induce slow-wave sleep (AP = −5.2, ML = ±0.7 mm from Bregma, DV = −4.2 mm from the skull). In a subsequent surgery, a craniotomy in the left hemisphere over the target site (AP = −2.0 mm, ML = −1.25 mm from Bregma) were made and covered with Kwik-Cast sealant (WPI). For mice used for the slow-wave sleep analysis, an additional 0.4 mm diameter catheter was implanted under the mice’s upper back skin to deliver clozapine N-oxide (CNO)^45^.

### Histology and probe position determination

Mice were transcardially perfused with 4% paraformaldehyde (PFA) solution and the extracted brains were stored in 4% PFA. 48 hours before sectioning, brains were transferred into 30% sucrose solution (diluted in 1X phosphate-buffered saline) and 50 μm or 80 μm coronal section were cut throughout the entire hippocampus. For the freely moving experiments, slices were Nissl-stained to visualize cell layer and probe tracks. For the head-fixed experiments, the tips of the silicon probes were coated with fluorescent DiD dye (Invitrogen) and the slices were stained with DAPI (Invitrogen). In both cases, probe placement was further verified by identifying the laminar layers of DG based on the CSD profile of dentate spikes.

### Recording sessions and analyzed data

In the freely moving experiments, electrophysiological recordings were performed at 30 kHz sampling rate, utilizing either ONIX^61^ or an OpenEphys Acquisition Board (OpenEphys). Camera and electrophysiology were synchronized in a custom Bonsai RX workflow. Freely moving QW sessions were recorded in a circular sleeping box (14 cm diameter) in which mice were allowed to move around (data A). Freely moving RUN sessions were recorded in an open-field arena in which mice were exposed to external sensory stimuli (visual, audio) and received a 5% sucrose drop as reward during the behavioral task. Analysis was restricted to epochs during high-speed locomotion (> 5 cm/s).

In the head-fixed experiments, prior to recordings, a 256-channel silicon probe (UCLA, 25 μm vertical spacing) was lowered through the craniotomy at a speed of 1 μm/s targeting the dorsal hippocampus, and a settling period of 45-60 min was ensured before data acquisition. Data was acquired at 30 kHz sampling rate utilizing an Intan acquisition system (Intan Technologies, CA, USA). In the head-fixed QW sessions (data B), mice were head-fixed on a spherical running ball (80 cm circumference) on an axle. Mice were allowed to run on the ball during the recordings but did not receive external sensory stimuli. The head-fixed RUN sessions were recorded during a virtual reality task on the spherical running ball in which mice received liquid milk reward. Speed of the mice were measured by utilizing a 512-bit rotary encoder (E5 optical kit encoder, US Digital) that was connected to the axle and analysis was restricted to running periods (> 5 cm/s). Electrophysiological and behavioral data were synchronized in the Intan acquisition system. In a separate head-fixed cohort, SWS was induced and maintained for several hours by injecting CNO via a catheter to activate GABAergic interneurons in the parafacial zone (0.5 mg/kg, injections in 3 h intervals)^45,62^. Data of the first three hours were analyzed in this study.

An additional dataset (data C) with pre-detected dentate spikes was analyzed that was downloaded from https://data.mrc.ox.ac.uk/waveform. Recordings were performed during quiet wakefulness and for a subset of mice the sessions were restricted to immobility^40^.

### Dentate spike classification

For recordings with double-shank probes, one shank was selected and used for analysis. In the head-fixed experiments, the shank layout consisted of 3 columns with 20 horizontal μm spacing, therefore only two neighboring columns were selected and their horizontal distance was ignored during the analysis. LFP was downsampled to 1-2 kHz, bandpass filtered in the 10-200 Hz range and broken channels were interpolated. Note that channels were not corrected to impedance, but approximately equal values were assumed. High impedance channels that were visually distinct from others were treated as broken channels and were interpolated. A channel located in the hilus was selected and putative dentate spikes with at least 4 standard deviations above mean were detected and width was restricted to the 5-50 ms range. Detection was repeated on a reference channel located outside the DG and events that coincided on the two channels (± 20 ms) were discarded. In data C, putative DS were pre-detected utilizing a different algorithm described in^40^. Current-source density around each putative DS was calculated and smoothed with a Gaussian filter across channels (σ = 1.5 channels). Next, PCA was performed on the CSD at each DS peak. We found that clustering DS in the space defined by the first two principal components was not always sufficient to separate the outer and middle molecular layers (oml, mml) from each other (k-means, Fig S1B). Therefore, two subsets of DS were selected: 1) events that exceeded the 75th percentile of the most negative first principal component values; and 2) events that exceeded the 75th percentile of the most positive first principal component values. The mean CSD of the two subsets were calculated, and the location of the deepest current sinks were determined. These two channels were representing the oml and mml respectively. DS were then projected into the oml-mml state space and events with a single sink in the oml or in the mml were considered as either DS1 or DS2. Next, DS with sinks in both layers were identified and the relative current sink was calculated as

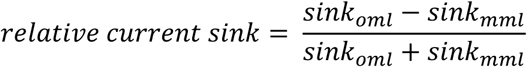

DS3 were defined as events with 1) current sinks both in oml and mml; and 2) relative current sink between −0.5 and 0.5. DS smaller than −0.5 relative current sink were considered as DS2 and events greater than 0.5 as DS1. The algorithms were implemented in a graphical user interface in MATLAB 2024b, allowing visual inspection and user-friendly application of the DS classification pipeline (https://github.com/EwellNeuroLab/DentateSpikeClassification).

### Speed analysis

In the freely moving experiments, behavior was recorded with a web camera (TECKNET 1080p Webcam) on a frame rate of 30 fps. The position of the body was post hoc tracked in DeepLabCut^63^ and instantaneous speed was calculated on the smoothed (167 ms wide moving average filter) position. In the head-fixed experiments, running speed measured by the rotary encoder was used. In both cases, running epochs were defined as a minimum of 0.5 s long continuous time window when mice exceeded 5 cm/s speed. Analyses for run-related DS were restricted to these running epochs.

### Single-unit analysis

Single-units were sorted in Kilosort1 or in Kilosort 2.5^64,65^ and then manually curated in Phy. Location of single-units were verified based on DS CSD profile or histology and cells in the DG were kept. Putative DG single units were analyzed in CellExplorer and classified as either principal cells or narrow waveform interneurons^66^. For each cell, peri-event time histograms were created in a ±200 ms time window around each dentate spike with a bin width of 1 ms, similar to^33^. A cell was considered active during an individual event, if it fired ± 10 ms from the DS peak. For measuring the proportion of neurons active during individual DS events, we excluded mice that had less than 10 cells recorded (total of 4 mice (15 cells) were excluded). Otherwise, every recorded neuron was included. Firing rate was averaged across DS events in each time bin and z-scored across time. The resulting trace was smoothened with a 3-point moving average filter. Maximum value in a ± 10 ms window around the DS peak was considered as the activity score of the cell. We considered a cell to be significantly active if the activity score exceeded z = 2.58, corresponding to 𝛼 = 0.01 significance level. To assess the influence of cell type (principal cell/interneuron), DS type (DS1/2/3) and brain state (QW/RUN) on cell recruitment, a linear mixed-effect model (LME) was constructed (*fitlme()*, MATLAB) to model the observed activity scores as

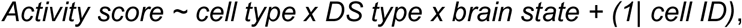

in which the intercept term (baseline: IN, DS1, QW), the three main effects and all their possible interaction terms were considered as fixed-effects. A random effect was included to account for repeated measurements within cells. The significance of individual fixed effects was assessed using ANOVA on the LME. Post hoc comparisons were performed using the *coefTest()* function in MATLAB by creating a contrast vector for each comparison respectively. P-values were then Bonferroni corrected for the overall 24 comparisons. Note that RUN and QW sessions were recorded subsequently, allowing the direct comparison of cells in the two brain states.

### Statistics

All statistical analysis was performed in MATLAB R2024b. Prior to statistical comparison, data was tested for normality (Anderson-Darling test).

### Data and code availability

Graphical user interface for the classification pipeline and additional codes used for the analysis are available on the Ewell lab GitHub page (https://github.com/EwellNeuroLab/DentateSpikeClassification).

## ACKNOWLEDGEMENTS

This work was funded by NIH R01 1R01NS128222-01 (to L.A.E.), Support Natural Sciences and Engineering Research Council of Canada (NSERC) Grant #RGPIN-2023-04402 (to B.L.M.), NIH Grant #NS132041 (to B.L.M.) and NIH R01 NS121764 (to B.L.M.).

## FIGURES AND TABLES

**Figure S1.**
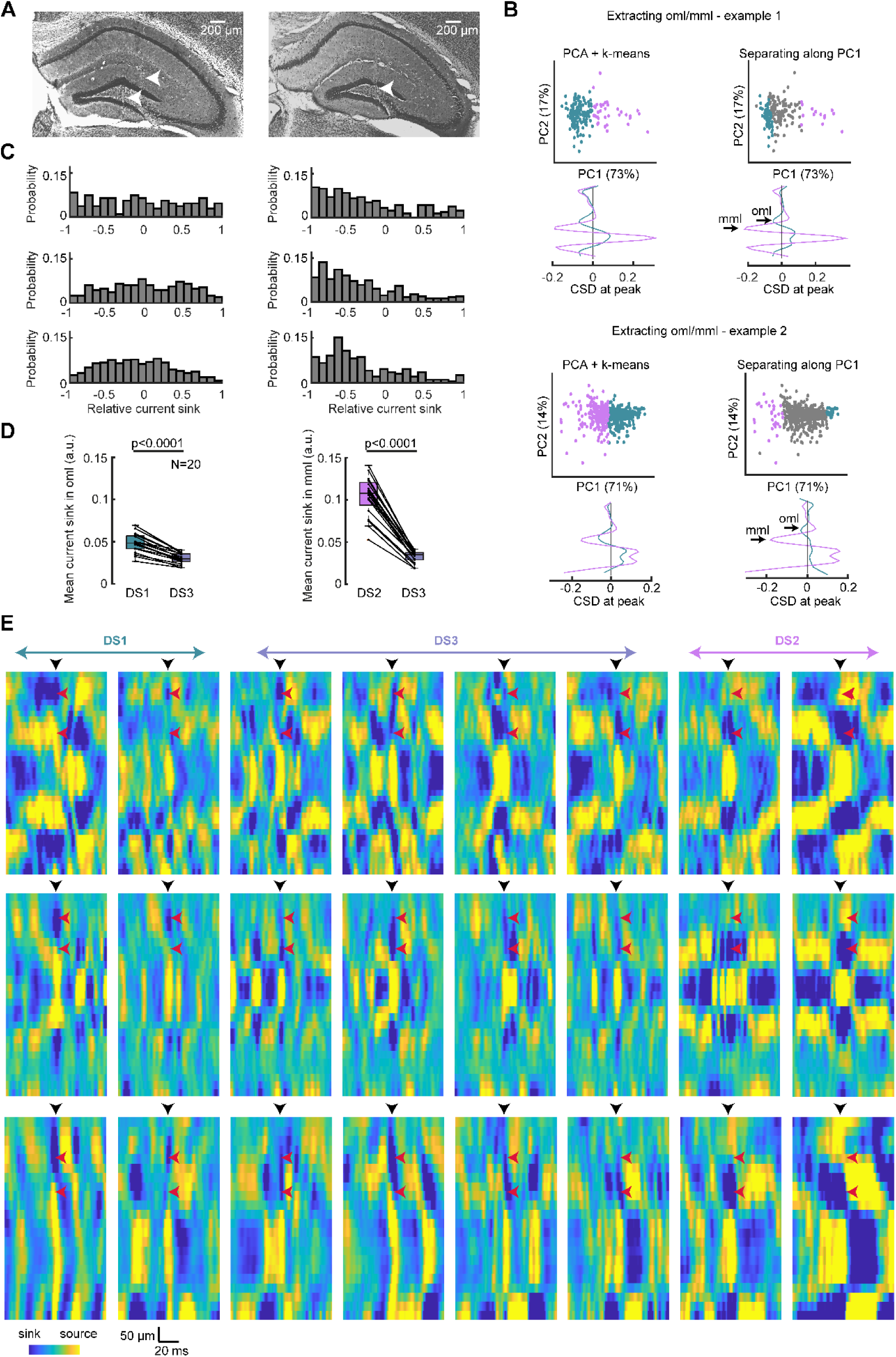
Multiple examples of DS in individual animals. (**A**) Examples of probe tracks in the DG in two different mice. (**B**) Two examples of oml/mml channel extraction. DS events were mapped into the space of the first two principal components. We found that k-means clustering in the space of the first two principal components was not always sufficient (example 2, left). Therefore, we used two subsets that exceeded the 75th percentile of most negative and most positive first principal components values (example 1-2, right). (**C**) Example of relative current sink distribution from six different mice. (**D**) Mean magnitude of current sinks in the outer molecular layer during DS1 and DS3 (right) and in the middle molecular layer during DS2 and DS3 (left). Mean magnitude was calculated for N = 20 mice. (**E**) Example of DS from three different mice. Black arrows mark the peak of the DS, red arrows indicate oml/mml respectively. Related to Figure 1.

**Figure S2.**
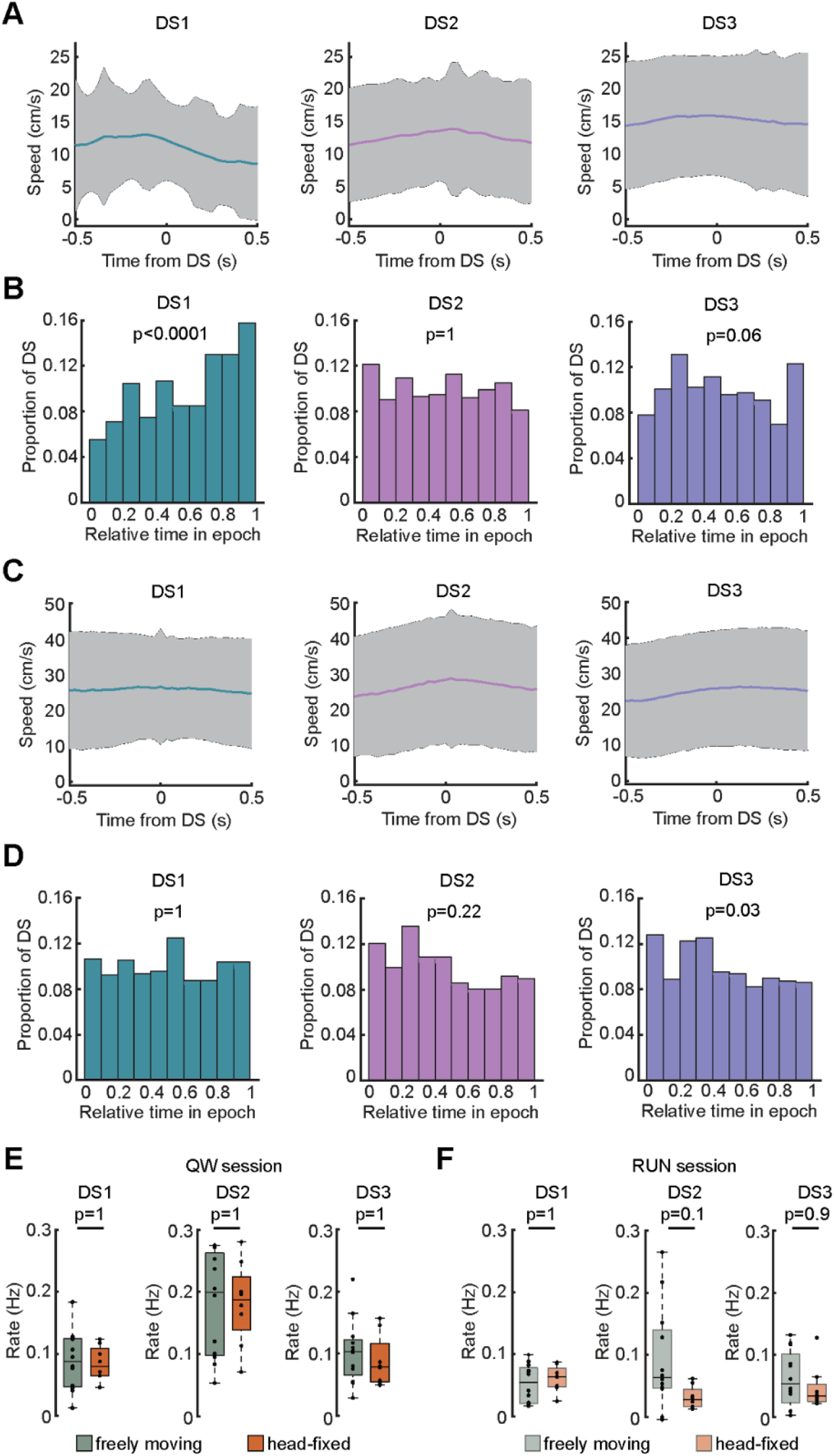
The influence of running speed and paradigm type on running-related DS. (**A**) Mean running speed before/after a dentate spike (colored lines) in freely moving mice. Gray shading marks standard deviation. (**B**) DS occurrence during individual running epochs for each DS type in freely moving mice. Note that DS1 proportion was higher at the end of the epochs, while DS2 and DS3 were distributed uniformly. (**C**) Same as in (A), but for head-fixed mice. (**D**) Same as in (B) but for head-fixed mice. (**E**) Comparison of DS rates between head-fixed and freely moving mice during QW. (**F**) Comparison of DS rates between head-fixed and freely moving mice during QW. Related to Figure 2.

**Figure S3.**
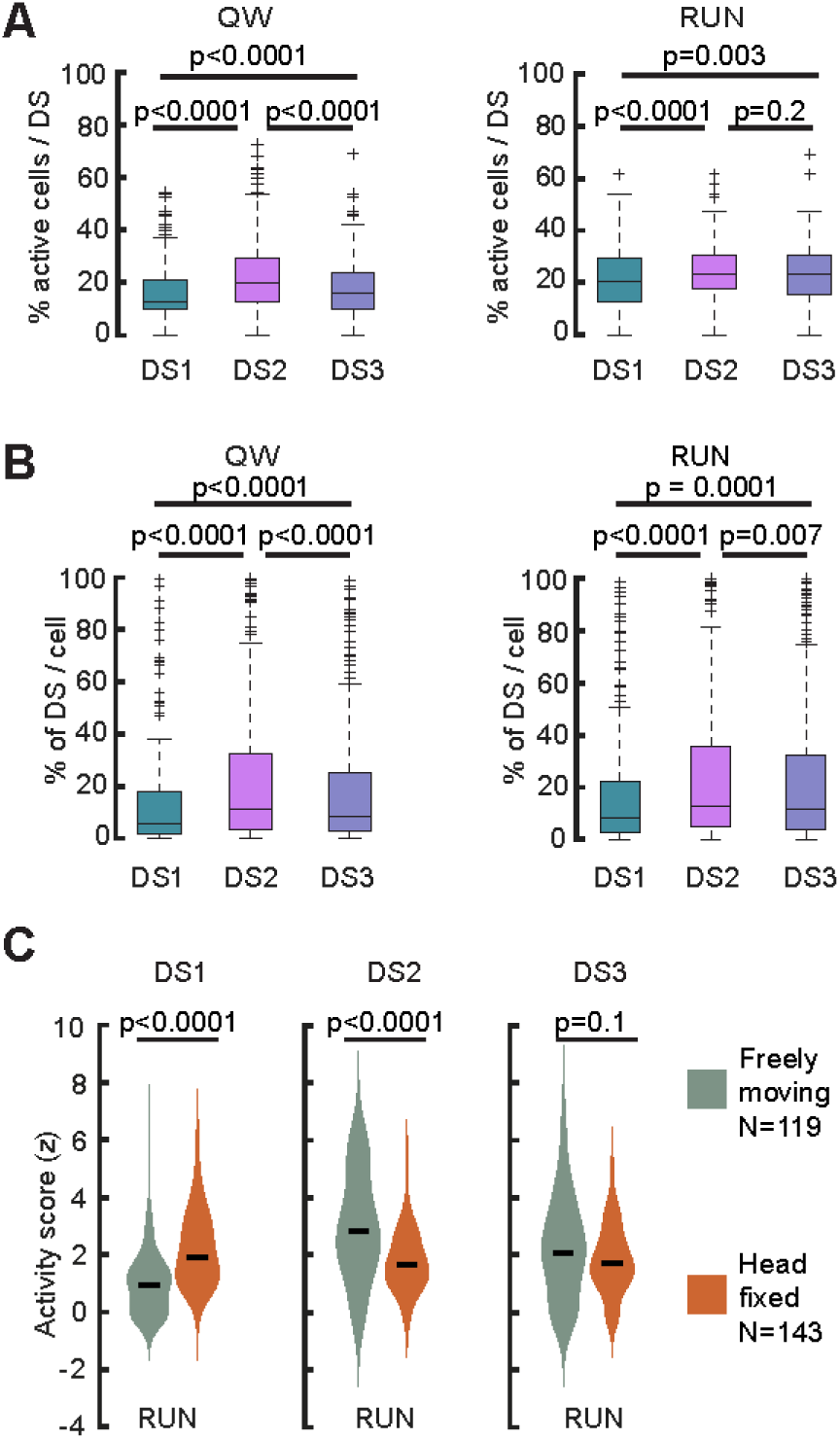
Cell activity and recruitment during DS. (**A**) Percentage of recruited neurons during individual DS during quiet wakefulness (left; DS1: N = 2384, DS2: N = 4947, and DS3: N = 2230) or running periods (right; DS1: N = 851, DS2: N = 345, DS3: N = 396). Cross markers indicate outliers. Mice with at least 10 neurons were included in this analysis. (**B**) Percentage of DS1, DS2 and DS3 during which individual neurons are active (N = 247). Cross markers indicate outliers. Mice with at least 10 neurons were included in this analysis. (**C**) Activity score of the freely moving (green) and head-fixed (orange) cell population for the three DS types during running. Black line marks the median of the distributions. Related to Figure 3.

**Supplementary Table 1.** Fitted linear mixed-effect model on the activity score of the recorded DG neurons. Related to Figure 3.

| Term | Estimate | Standard error | Degree of freedom | t-statistics | p-value |
| --- | --- | --- | --- | --- | --- |
| Intercept | 3.01 | 0.17 | 1560 | 17.68 | < 0.0001 |
| CellType_PC | -1.23 | 0.21 | 1560 | -5.92 | < 0.0001 |
| DSType_DS2 | 3.19 | 0.19 | 1560 | 16.4 | < 0.0001 |
| DSType_DS3 | 1.59 | 0.19 | 1560 | 8.16 | < 0.0001 |
| State_RUN | -0.95 | 0.19 | 1560 | -4.9 | < 0.0001 |
| PC:DS2 | -1.15 | 0.24 | 1560 | -4.86 | < 0.0001 |
| PC:DS3 | -0.7 | 0.24 | 1560 | -2.93 | 0.003 |
| PC:RUN | 0.62 | 0.24 | 1560 | 2.6 | 0.009 |
| DS2:RUN | -2.06 | 0.28 | 1560 | -7.49 | < 0.0001 |
| DS3:RUN | -0.98 | 0.28 | 1560 | -3.56 | 0.0004 |
| PC:DS2:RUN | 0.48 | 0.34 | 1560 | 1.43 | 0.15 |
| PC:DS3:RUN | 0.32 | 0.34 | 1560 | 0.95 | 0.34 |

